# *In vitro* activity of hydroxychloroquine in combination with antibiotics against intracellular uropathogenic *Escherichia coli*

**DOI:** 10.1101/2024.10.13.618068

**Authors:** V. Pargny, N. Vautrin, D. Ribet, O. Minlong, F. Caron, M. Pestel-Caron, K. Alexandre

**Author notes:** **Corresponding author** Kévin Alexandre, Hôpital Charles Nicolle, Centre Hospitalo-Universitaire, 37 Bd Gambetta, 76000 Rouen, France.

## Abstract

**Background:** Two pathophysiological concepts may explain recurrent UTI: a reinfection by a bacterial strain from the digestive microbiota or a relapse from *Escherichia coli* persisting within the superficial urothelial cells as intracellular bacterial communities (IBC). Hydroxychloroquine (HCQ)-antibiotic combination is effective against intracellular adherent-invasive *E. coli* isolated from patients with Crohn’s disease. We hypothesized that HCQ may enhanced antibiotic efficacy against *E. coli* IBC.

**Methods:** UTI89 reference strain and two clinical *E. coli* strains forming IBCs (VITALE#2157, VITALE#2206) were used in this study. MIC, MBC and time killing curves (4xMIC) were performed for AZYTHROMYCIN (AZT), CIPROFLOXACIN (CIP), DOXYCYCLINE (DC), FOSFMOMYCIN (FF) and HCQ. *In vitro* activity of antibiotics alone and combined with HCQ was evaluated on intracellular bacterial survival assay using a model of urothelium cells (HTB-9).

**Results:** Time-killing curves showed that CIP has a bactericidal activity at H6 (-4.6 log_10_ CFU/mL) with regrowth at H24 against 2157, and a bactericidal activity at H6 without regrowth at H24 against 2206 (-3.3 log_10_ CFU/mL) and UTI89 (-3.4 log_10_ CFU/mL); DC, AZT and FF demonstrated bacteriostatic activity at H24 regardless of the strain. We observed a significant decrease in the number of intracellular bacteria at H42 with CIP (-1.83 log_10_ / 10_6_ cells). DC, FF, and AZT exposure did not reduce IBC formation at H42 (P>0.05). IBC formation after HCQ-antibiotic combination exposure was not significantly different (P>0.05) from antibiotic exposure alone, regardless of the antibiotic or strain studied.

**Conclusions:** In conclusion, HCQ exposure does not enhance antibiotics activity against intracellular uropathogenic *E. coli*.

**Highlights:**

- Using an *in vitro* model of superficial urothelium monolayer, we observed a significant decrease in intracellular uropathogenic *Escherichia coli* in response to ciprofloxacin exposure.
- Hydroxychloroquine exposure does not enhance antibiotics activity against intracellular uropathogenic *Escherichia coli*.

## I. Introduction

Urinary tract infections (UTIs) are considered as the most common bacterial infections. Uropathogenic *Escherichia coli* (UPEC) strains are involved in 70 to 95% of cystitis in women, the most common presentation of UTI.^1^ Recurrent cystitis (RC) is defined as the occurrence of at least four episodes of cystitis over 12 months,^2,3^ and affects about 5% of women. Currently, besides the correction of contributing factors, the main recommended therapeutic alternatives are iterative or continuous antibiotics treatment.^3^

Two pathophysiological concepts may explain RC: (i) a reinfection by a bacterial strain from the digestive microbiota;^4^ (ii) a relapse from UPEC persisting within the urothelium. Indeed, UPECs are able to constitute intracellular bacterial communities (IBCs) within superficial urothelial cells, and quiescent intracellular reservoir (QIR) within transitional urothelial layer cells.^1,5^ Antibiotic mono-therapy does not show *in vivo* activity against IBC or QIR.^6,7^

Hydroxychloroquine (HCQ) is a lysosomotropic agent that increases endosomal pH, thus inhibiting lysosomal enzymes and blocking autophagy.^8^ Doxycycline-HCQ combination is effective against intracellular *Coxiella burnetti*.^9^ This combination is actually the first line treatment recommended for *C. burnetti* and *Trophyrema whipplei* endocarditis.^10^ Some data suggests that HCQ could also promote the elimination of intracellular *E. coli* within Crohn’s disease monocyte-derived macrophages.^11^

Therefore, we aimed to study the *in vitro* efficacy of HCQ in combination with antibiotics against intracellular UPEC.

## II. Material and methods

### II.1. Bacteria and bladder cell culture

Two clinical strains (VITALE#2157, VITALE#2206) from a prospective study (Vitale study, ClinicalTrials.gov identifier: NCT02292160) and collected from urine of women suffering from community-acquired RC were selected and characterized as forming IBCs. UTI89 (acute cystitis) was used as a UPEC IBC producer reference strain. Al l strains were stored in glycerol medium at -80°C. Strains were thawed as needed on agar (Luria-Bertani agar LBA, Sigma-Aldrich; or tryptone soy agar TSA, Bio-Rad) or in Luria-Bertani broth.

Bladder cell culture was performed using the human epithelial cell line HTB-9 (ATCC 5637). This cell line is derived from a urothelial carcinoma and is able to form a superficial unistratified epithelium.^6^ Cell cultures were maintained at 37°C and 5% CO_2_ in RPMI 1640 medium (Gibco® A10491, ThermoFischer-Scientific) supplemented with 10% fetal bovine serum.

### II.2. Molecules and Antibiotic Susceptibility Testing

All antibiotics were purchased from Sigma-Aldrich and used at the final concentrations indicated. Stocks were prepared as follows: fosfomycin (FF) 10 g/L, ciprofloxacin (CIP) 10 g/L, doxycycline (DC) 10 g/L, azithromycin (AZT) 10 g/L and hydroxychloroquine (HCQ) 2.5 g/L. CIP was solubilized in a 0.1 mmol/L HCl aqueous solution. AZT was solubilized in 100% ethanol.

Minimum inhibitory concentrations (MICs) were measured by broth microdilution (BMD) for AZT, CIP, DC and HCQ according to CLSI (CLSI-M07-A9), except for HCQ (adapted from CLSI). FF MIC was measured by agar dilution with a G6PD supplemented media as recommended (CLSI-M07-A9). All experiment were performed in triplicate. Quality control was performed with the *E. coli* ATCC 25922 strain on each batch.

### II.3. Bactericidal activity assays

Minimal bactericidal concentrations (MBCs) were evaluated by BMD for AZT, CIP, FF and DC in triplicate according to CLSI recommendations (CLSI-M07-A9). Concentration ranged from 0.5 to 16 x MIC.

Time-killing curves were performed for each antibiotic (excepted for HCQ). Final antibiotic concentration was 4 x MIC. Mueller Hinton broth (MHB) used for FF was supplemented with glucose-6-phosphate (25 mg/L), as recommended by CLSI. Lower limit of detection was 1.3 log_10_ CFU/mL. All experiment were performed in duplicate.

### II.4. Cytotoxicity

The cytotoxicity of antibiotics and HCQ towards HTB-9 cells was evaluated using the CytoTox 96® Non-Radioactive Cytotoxicity Assay (Promega). This test was performed in triplicate for each antibiotic alone (AZT 1 and 20 mg/L; CIP 50 and 100 mg/L; FF 100 and 1000 mg/L; DC 10 and 50 mg/L) or in combination with HCQ (HCQ 10 mg/L), following supplier’s protocol.

### II.5. Intracellular bacterial survival assays

Internalization of VITALE#2206, VITALE#2157 and UTI89 strains into HTB-9 cells, as well as the impact of antibiotics and HCQ on intracellular survival were evaluated as previously described.^6^ Briefly, strains were thawed on chromogenic agar (CPSE, bioMérieux) and were grown at 37°C for 48 h in static LB broth to induce type 1 pilus expression.^12^ Triplicate sets of confluent HTB-9 bladder epithelial cell monolayers grown in 12-well tissue culture plates (with a density of 2.5 x 10^5^ cells/cm^2^) were infected with UPEC strains using a multiplicity of infection (MOI) of 15 bacteria per host cell. HTB-9 cells were further incubated for 24 h to enable intracellular bacteria growth. After 24 h, cells were washed twice and incubated for 18h in culture medium containing gentamicin (10 mg/L) and the different antibiotics studied, with or without HCQ. Control wells (enumerated at H24 and H42 post-inoculation) were exposed to culture-medium without the studied antibiotics. Cells were lysed 42 h post-inoculation and cell lysates were plated on LB agar plates to enumerate viable intracellular bacteria.

### II.6. Statistics

Bacterial counts before and after HCQ and/or antibiotic exposure were described as median (interquartile range [IQR]) and compared by Wilcoxon test. Data analysis and plots were performed with R software (R Foundation for Statistical Computing, Vienna, Austria, https://www.R-project.org/).

## III. Results

The origin and IBC formation capacity of strains as well as MIC and MBC of antibiotic against each strain are described in supplementary table 1. As expected HCQ denoted no antibacterial activity alone (MIC>10 mg/L). The internalization rate (*i*.*e*., the ratio between intracellular bacteria and the number of bacteria added to cells) varied between strains from 0.03% to 0.11%. The number of intracellular bacteria at H42 varied accordingly between 3.11 to 5.04 log_10_ CFU / 10^6^ cells.

Figure 1 shows time-killing curves according to strains. CIP was rapidly bactericidal (as early as H6) against all strains. Nevertheless, at H24 a regrowth was observed for CIP against the VITALE#2157 strain. AZT and DC showed bacteriostatic effect against all strains. FF demonstrated a fast (H3 to H6) and significant (∼3 log_10_ CFU/mL) reduction of viable bacterial population but with a regrowth of all strains at H24.

**Figure 1:**
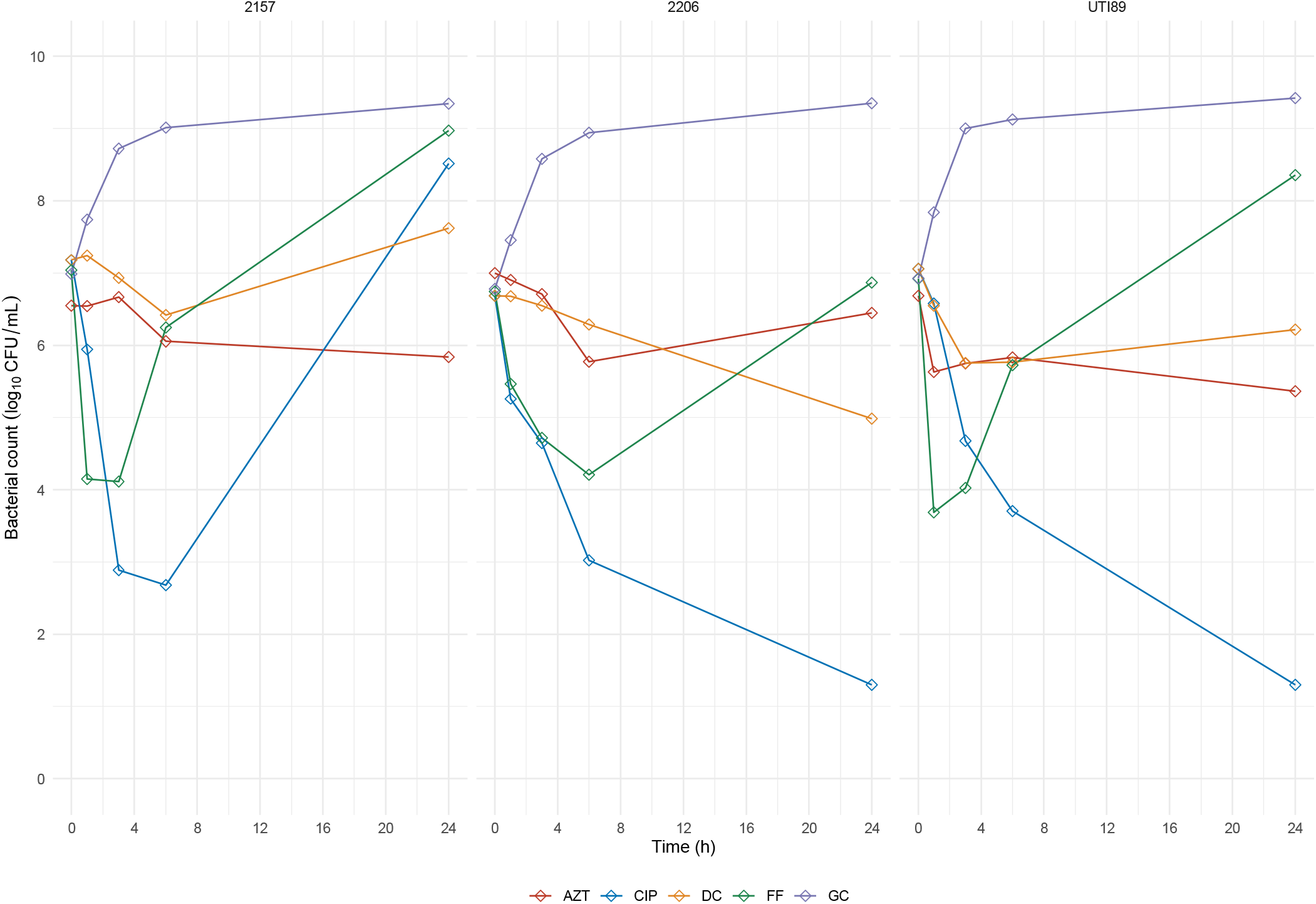
Time killing assays according to strains and antibiotics. All antibiotics were tested at a concentration equivalent to 4 x MIC for each strain. AZT: azithromycin, CIP: ciprofloxacin, DC: doxycycline, FF: fosfomycin, GC: growth control. Dots and lines represent mean of assays performed in duplicate. The lower limit of detection was 1.3log_10_ CFU/mL.

All the tested molecules (except DC at 50 mg/L concentration) were considered as non-cytotoxic (*i*.*e*., cytotoxicity level was lower than or equal to the basal level of cell death without antibiotics, Supplementary Table 2).

As showed in Figure 2, compared to untreated control, we observed a significant decrease in the number of intracellular bacteria compared to untreated control at H42 with CIP 2 mg/L: -1.83 (IQR=0.67) log_10_ / 10^6^ cells (p<0.001) and with CIP 100 mg/l: -1.79 (IQR=1.34) log_10_ / 10^6^ cells (p=0.004). AZT 20 mg/L, DC 10 mg/L, FF 100 mg/L and FF 500 mg/L exposure did not impair the number of intracellular bacteria at H42 compared to controls: -0.55 (IQR=0.67) log_10_ / 10^6^ cells (p=0.387), -0.30 (IQR=0.30) log_10_ / 10^6^ cells (p=0.401), -0.37 (IQR=0.82) log_10_ / 10^6^ cells (p=0.094), -0.43 (IQR=0.93) log_10_ / 10^6^ cells (p=0.077) respectively. The number of intracellular bacteria after HCQ-antibiotic combination exposure was not significantly different from antibiotic exposure alone, regardless of the evaluated antibiotic agent or bacterial strain (Figure 2).

**Figure 2:**
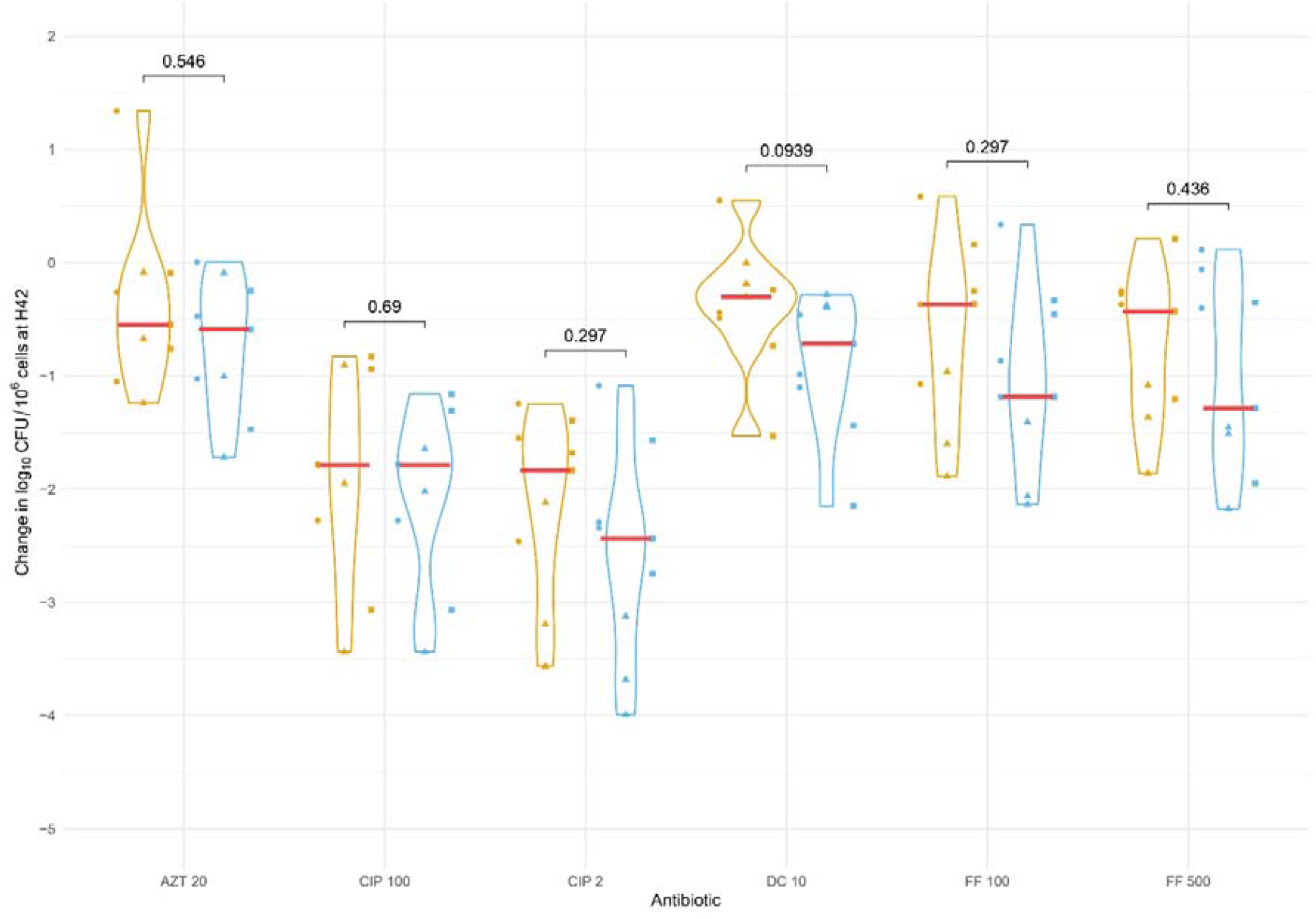
Violin plots of change in the number of intracellular bacteria (log_10_ CFU per 10^6^ cells) compared to controls at H42 post-infection, according to antibiotics and HCQ combination. IBC: intracellular bacterial communities. HCQ: hydroxychloroquine. Orange: antibiotic alone. Blue: HCQ-antibiotic combination. Circle dot: 2157 strain. Triangle dot: 2206 strain. Square dot: UTI89 strain. AZT 20: azithromycin 20 mg/L. CIP 100: ciprofloxacin 100 mg/L. CIP 2: ciprofloxacin 2 mg/L. DC 10: doxycycline 10 mg/L. FF 100: fosfomycin 100 mg/L. FF 500: fosfomycin 500 mg/L. Red line: IBC median change. P-value above bracket were calculated between antibiotic alone and HCQ-antibiotic combination group by Wilcoxon test.

## IV. Discussion

In an *in vitro* unistratified urothelium model, we observed no significant difference in the decrease of intracellular uropathogenic *E. coli* after treatment with antibiotics alone or in combination with HCQ.

Several data may explain our results. First, HCQ is a lysosomotropic agent able to accumulate in acid vesicles and increase their pH.^8^ Previous study have shown that replication of adherent invasive *E. coli* within macrophage phagolysosomes was significantly decreased by HCQ.^11^ Nevertheless, it appears that within urothelial superficial cells, UPEC are internalized in fusiform-shaped vesicles (Rab27b^+^) before exiting into the cytosol in order to form IBCs. UPEC-containing vesicles in superficial cells do not fuse with lysosomes and bacteria rapidly exit from these vesicles (as early as 1h post-infection).^13^ In our model, we treated urothelial cells 24-h post-infection. UPEC are at this stage probably already within the cytosol forming IBCs. This could explain the inefficiency of HCQ.

Second, in transitional cells, UPEC are internalized and form QIR in late acidic endosomes.^14^ Although the regulation mechanisms of this quiescent state are still not well known, it could be assumed that HCQ will disrupt the internalized UPEC metabolism by inhibiting endosomal acidification. Nevertheless, our monolayer superficial urothelium model did not allow us to study the potential HCQ activity on QIR established in transitional urothelial layer.

Our results on CIP efficacy against intracellular uropathogenic bacteria stress the importance of studying QIR for RC pathophysiology. Indeed, these results are in accordance with the well-known fluoroquinolones activity on intracellular bacteria,^15^ and with previous work showing that fluoroquinolones are efficient against IBCs in superficial urothelium models.^6,7^ Of note, Blango *et al*. did not observe any decrease in bacterial count within bladder tissue after fluoroquinolones exposure in a murine model of chronic cystitis.^6^ These results can be explained by the presence of QIR within the transitional cells.

In conclusion, HCQ exposure does not promote fluoroquinolones bactericidal activity against intracellular uropathogenic bacteria. Further work is now needed to assess the potential efficacy of HCQ-antibiotics combination against QIR.

## Ethical approval

Not required

## Sequence information

Not applicable

## VIII. Appendice

**Supplementary Table 1:** Strains main characteristics, MICs and MBCs according to antibiotic.

| Characteristics and *in vitro* outcomes | Strains |  |  |
| --- | --- | --- | --- |
|  | VITALE#2157 | VITALE#2206 | UTI89 |
| **Characteristics** |  |  |  |
| Origin | VITALE study | VITALE Study | IAME UMR 1137 |
| Type of UTI | recurrent cystitis | recurrent cystitis | acute cystitis |
| Phylogroup | A | B2 | B2 |
| **IBC** |  |  |  |
| Mean (SD) internalization rate (%) | 0.0303 (7.4e-3) | 0.1148 (8.96e-2) | 0.0338 (3.00e-4) |
| Median (IQR) log<sub>10</sub> CFU / 10<sup>6</sup> cells H42 | 3.11 (1.55) | 5.04 (0.90) | 3.98 (0.51) |
| **MIC (mg/L)** |  |  |  |
| Azithromycin | 8 | 4 | 8 |
| Ciprofloxacin | 0.008 | 0.008 | 0.008 |
| Doxycycline | 0.5 | 0.5 | 1 |
| Fosfomycin | 1 | 1 | 2 |
| Hydroxychloroquine | >10 | >10 | >10 |
| **MBC (mg/L) – (n x MIC)** |  |  |  |
| Azithromycin | >128 (>16) | >64 (>16) | >128 (>16) |
| Ciprofloxacin | 0.016 (2) | 0.008 (1) | 0.016 (2) |
| Doxycycline | >8 (>16) | >8 (>16) | >8 (>16) |
| Fosfomycin | 2 (2) | 2 (2) | 4 (2) |
| Hydroxychloroquine | NA | NA | NA |

MIC: minimum inhibitory concentration, MBC: minimum bactericidal concentration, AZT: azithromycin, CIP: ciprofloxacin, DC: doxycycline, FF: fosfomycin

**Supplementary Table 2:** Cytotoxic effects of antibiotics and HCQ combinations on HTB-9 cells. NC, no-cell (culture medium only); VOC, vehicle only cell (cells and medium without antibiotics); MLR, maximum LDH release (cells and medium without antibiotics, after saponin lysis).

| **Conditions** | **LDH release (% of MLR)** | **Combination**<br>**(ATB and HCQ 10 mg/L)**<br>**LDH release (% of MLR)** |
| --- | --- | --- |
| NC | 0 | NA |
| VOC | 26 | NA |
| MLR | 100 | NA |
| HCQ (10 mg/L) | 18 | NA |
| FF (1000 mg/L) | 31 | 16 |
| FF (100 mg/L) | 20 | 25 |
| AZT (20 mg/L) | 24 | 26 |
| AZT (1 mg/L) | 29 | 18 |
| CIP (100 mg/L) | 28 | 10 |
| CIP (50 mg/L) | 15 | 15 |
| DC (50 mg/L) | 44 | 49 |
| DC (10 mg/L) | 20 | 20 |

